# Multicellular feedback control of a genetic toggle-switch in microbial consortia

**DOI:** 10.1101/2020.03.05.979138

**Authors:** Davide Fiore, Davide Salzano, Enric Cristòbal-Cóppulo, Josep M. Olm, Mario di Bernardo

## Abstract

We describe a multicellular approach to control a target cell population endowed with a bistable toggle-switch. The idea is to engineer a synthetic microbial consortium consisting of three different cell populations. In such a consortium, two populations, the Togglers, responding to some reference input, can induce the switch of a bistable memory mechanism in a third population, the Targets, so as to activate or deactivate some additional functionalities in the cells. Communication among the three populations is established by orthogonal quorum sensing molecules that are used to close a feedback control loop across the populations. The control design is validated via in-silico experiments in BSim, a realistic agent-based simulator of bacterial populations.

## INTRODUCTION

The main goal of Synthetic Biology is the design of reliable genetic circuits able to endow living cells with new functionalities [1]. Examples include the development of bacteria able to sense and degrade pollutants (like hydrocarbons or plastic) in the environment [2], or cells that can track and kill cancer cells by releasing drugs at specific locations, limiting dangerous side effects [3].

Biological systems capable of carrying out these complex tasks require the integration of advanced functional components analogous to those of an autonomous robotic systems [4]. Specifically, sensors are needed to perceive stimuli from the environment, actuators to interact with the environment (e.g. production and delivery of desired molecules or drugs), and more importantly, some control logic with a memory mechanism able to make decisions and regulate the cell behavior. However, due to current technological and biological limitations, e.g. metabolic burden and retroactivity, it is hard to implement the entire control system inside a single cell [5]. A promising solution to overcome this problem is to assign the required functionalities to different cell populations such that each of them carries out a specific task [6], [7]. For example, one population can be specialized to sense a certain molecule in the environment and to communicate its presence to the rest of the consortium by secreting signaling molecules in its surroundings. In this way, more complex tasks can be carried out by multicellular systems as the result of the mutual interactions between their components [6].

In this letter we present a novel multicellular feedback control strategy involving a microbial consortium consisting of three cellular populations, in which the activity of one of them is governed by the other two. Specifically, the state of a genetic toggle-switch endowed in one of the populations, the *Targets*, can be controlled by providing or removing a reference input to the other two, the *Togglers*, which communicate with the Targets via orthogonal quorum sensing molecules. In this way, it can be possible to toggle additional functionalities in the Targets, as required in a number of applications, e.g. production and secretion of some desired molecule or drug in the environment [3], [8].

The relationship between the three cell populations in the consortium and their molecular signals can be schematically represented as a sequential logic circuit [9] (Fig. 1). The two controller cells sense the concentration in the environment of the reference signal Ref and of the Targets’ output *y*, which is high (*y* =1) only when the Targets are active. The controllers then generate two control signals *u*_1_ and *u*_2_ according to the following logic functions

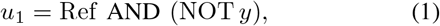

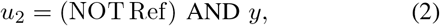

so that the reference signal, Ref, can be used to toggle the switch between the ON state and the OFF state.

**Fig. 1.**
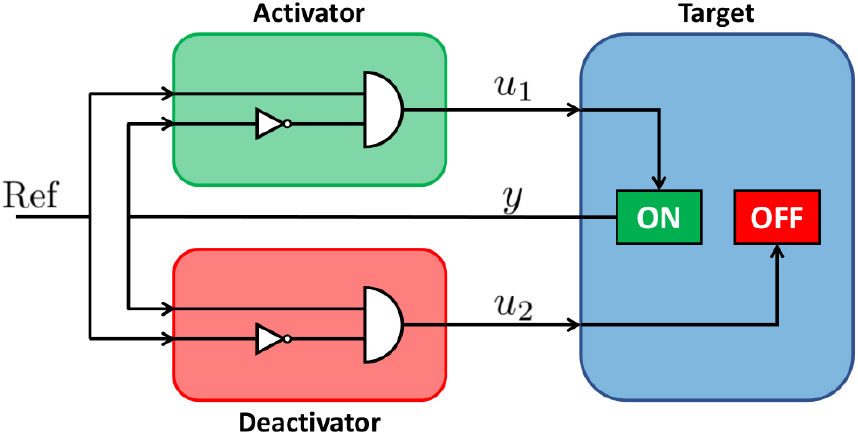
Representation as a sequential logic circuit of the relationship between the cell populations and their molecular signals. The Target receives ON commands (*u*_1_ = 1) only when Ref = 1 AND *y* = 0, and OFF commands (*u*_2_ = 1) only when Ref = 0 AND *y* = 1. In this way, the input signals *u*_1_ and *u*_2_ are equal to 1 only when there is disagreement between Ref and *y*.

In particular, a controller population, the *Activators*, command the activation of the Targets when, at the same time, they perceive the presence of a specific *reference* chemical signal in the environment and the Targets are inactive, while the other controller population, the *Deactivators*, inhibit the activity of the Targets when they are active and the reference signal is no longer present in the environment. In this way, the Targets are active only when the reference signal is present in the environment.

The crucial challenge we address in this letter is the biological implementation of this novel multicellular control scheme. After proposing a possible realization of all the functions required, we model the three populations and investigate analytically how to engineer the consortium parameters so as to guarantee its desired operation. We then provide in-silico experiments in a realistic agent-based simulator of bacterial populations confirming the viability of the approach.

## II. MULTICELLULAR CONTROL SYSTEM

A schematic biological implementation of the multicellular control strategy we propose is illustrated in Fig. 2. The superscripts e, t, a, d are used in the rest of the paper to denote quantities in the environment, Target cells, Activator cells and Deactivator cells, respectively.

**Fig. 2.**
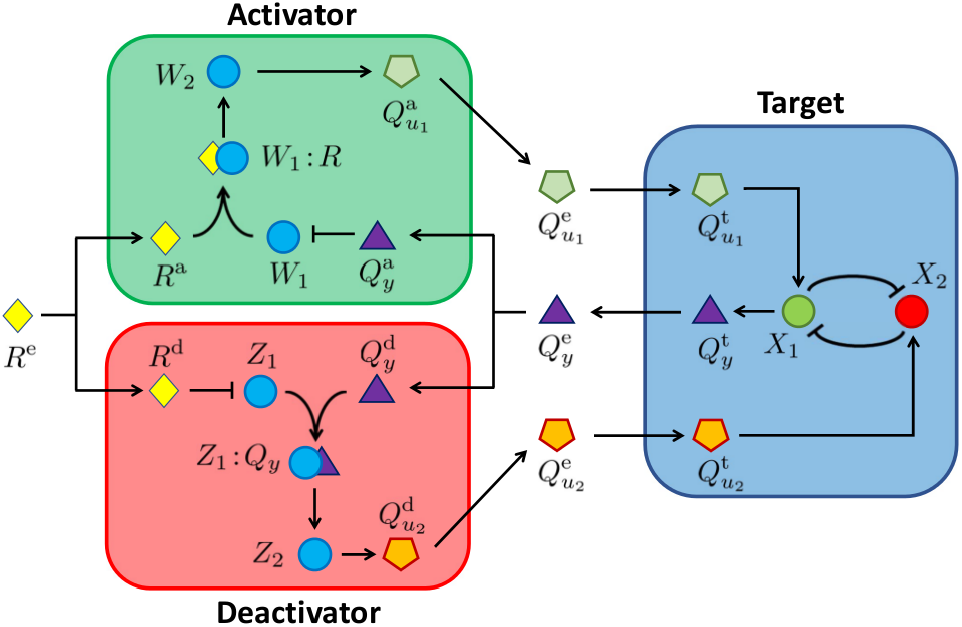
Abstract biological implementation of the multicellular control of a genetic toggle-switch. The Togglers compare the concentrations of the signaling molecules *R* and *Q_y_* using an antithetic motif and produce *Q_u_i__* according to the logic functions (1)-(2). This, in turn, diffuses inside the Target and promotes the activation of *X_i_*, which makes the Target change its state. Circles represent internal molecular species and polygons represent signaling molecules diffusing in the cells.

Activation and repression of each species is governed by Hill functions with dissociation coefficient *θ* and exponent *n*. Moreover, we denote with 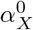 and *α_X_* the basal and maximal expression rates of species *X*, and with the *γ^υ^* degradation rate of a species within a domain *υ*.

### A. Target population

We assume that the bistable memory regulating the activation of the Target cells is implemented by an inducible genetic *toggle-switch* [10], [11]. This genetic network consists of two proteins, *X*_1_ and *X*_2_, each repressing the expression of the other, so that at steady state only one is fully expressed. Without loss of generality, we assume full expression of *X*_1_ corresponds to the “active” state of the cell where some desired functionalities are turned on, while full expression of *X*_2_ corresponds to its inactive state.

We focus here on the problem of toggling the Target population between the two states. Other works in the literature have considered the alternative control problem of stabilizing the toggle-switch about some intermediate expression levels of *X*_1_ and *X*_2_, see e.g. [11]–[15]; a problem we do not address in this paper.

The dynamical model of the toggle-switch can be given as

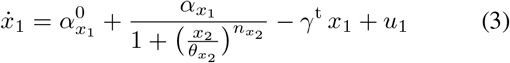

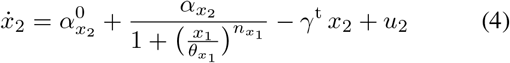

where the state variables *x*_1_ and *x*_2_ denote concentrations of molecules *X*_1_ and *X*_2_ inside the cell and we assume *u*_1_ and *u*_2_ capture the effect of two inputs that can be used to toggle the switch between one state and the other.

We assume that the parameters of the toggle-switch are chosen such that in the absence of external inputs, i.e. *u*_1_ = *u*_2_ = 0, the system is bistable [16], with well separated equilibria and sufficient transversality of the nullclines [17]. Specifically, system (3)-(4) admits two stable equilibria, 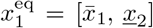 and 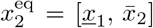, associated to high expression of species *X*_1_ or *X*_2_, respectively. We also assume, that there exists some positive value 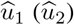 such that, when 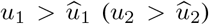 and *u*_2_ = 0 (*u*_1_ = 0), system (3)-(4) converges to a unique equilibrium point corresponding to high expression of *X*_1_ (*X*_2_) and remains therein when the inputs are switched off.

As shown in Fig. 2, we associate each of the inputs of the toggle-switch (3)-(4) in the Targets to the concentration of a quorum sensing molecule coming from the Activator and Deactivator cells. Specifically, we capture the promoting action of the signaling molecule *Q_u_i__* on the expression of *X_i_* by setting

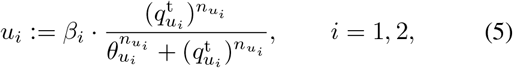

where 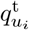 denotes the concentration of molecule *Q_u_i__* inside the Target cell, and *β_i_, θ_u_i__* and *n_u_i__* are the maximal promoter activity, activation and Hill coefficients, respectively.

In our design, Target cells can signal their state to the other cells by means of another, orthogonal, quorum sensing molecule *Q_y_* that is produced at rate 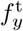 assumed to be proportional to *X*_1_ [7], that is

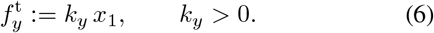

Hence, at steady state, when the cell is active, 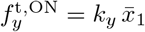.

### B. Toggler populations

The two controllers implement the same logic circuits (Fig. 1) and therefore they also share similar biological implementations. However, since the reference molecule *R* and signaling molecule *Q_y_* have inverted roles, the biochemical reactions describing the Activator and Deactivator cells are in general different. For the sake of brevity, we next describe only the biological implementation of the Deactivator cells in Fig. 2, which is directly taken from [7].

The logic function (2) is realized in the Deactivators by means of an antithetic motif. Specifically, the expression of *Z*_1_ is regulated by two independent and competing species, *R* and *Q_y_. R* represses *Z*_1_, while *Q_y_* activates *Z*_1_ and reacts with it forming the complex *Z*_1_: *Q_y_. Q*_*u*_2__ is then produced through a synthesis process catalyzed by *Z*_2_, which is promoted only by the active compound *Z*_1_: *Q_y_*. Asa result, the control signal molecule *Q*_*u*_2__ is produced and released only when the concentration of *R* inside the Deactivator cells is low while that of *Q_y_* is high.

By denoting with *z*_1_ and *z*_2_ the concentrations of the species *Z*_1_: *Q_y_* and *Z*_2_ in the Deactivators, their dynamics can be written as

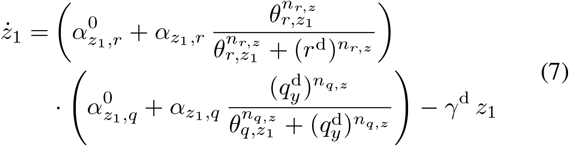

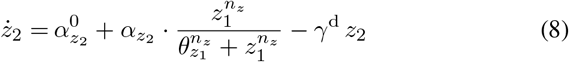

The output signaling molecule *Q*_*u*_2__ is produced through a synthesis process catalyzed by *Z*_2_, and so at a rate 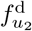 proportional to the concentration of *Z*_2_, that is

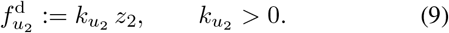

Moreover, the substrates required to synthesize *Q*_*u*_2__ are assumed to be in excess and therefore this process does not directly affect *Z*_2_ [7].

Similarly, another antithetic motif is embedded in the Activators so that by denoting with *w*_1_ and *w*_2_ the concentrations of the species *W*_1_: *R* and *W*_2_ therein, their dynamics can be written as

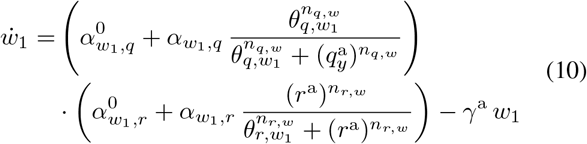

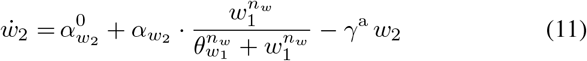

The Activators will then generate a quorum sensing molecule *Q*_*u*_1__ at a rate 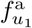 proportional to the concentration of *W*_2_, that is

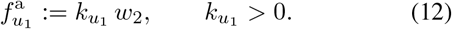

### C. Intercellular communication

The intercellular communication between the three populations is realized by means of three pairwise orthogonal quorum sensing molecules. Namely, *Q*_*u*_1__, *Q*_*u*_2__ and *Q_y_*, which are produced by Activators, Deactivators and Targets, respectively. For the sake of brevity, in what follows we use the placeholder superscript j to denote concentrations of signaling molecules in a generic cell type, where j = a for Activators, j = d for Deactivators and j = t for Targets. The quorum sensing molecules and the reference signal molecule *R* diffuse across the cell membrane of the genetic cell of type j with diffusion rate *η*^j^. The evolution of the concentrations of the signaling molecules *inside* the generic cell of type j can then be given as

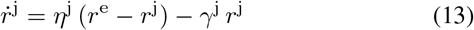

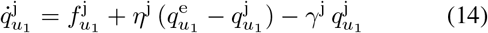

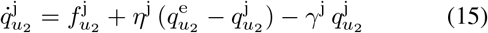

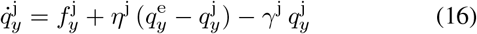

where the production functions 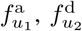 and 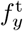 are defined in (12), (9) and (6), and 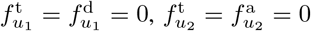. and 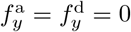.

The concentrations of the reference signal molecule and of the quorum sensing molecules secreted by the three cell populations in the environment can be described by the following set of ODEs:

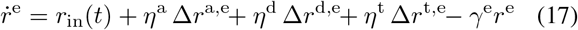

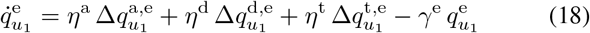

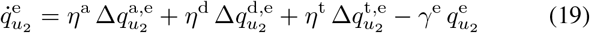

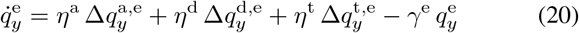

where Δ*r*^j,e^: = *r*^j^ − *r*^e^, j ∈ {a, d, t}, and similarly are defined 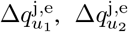, and 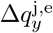, and the function *r*_in_(*t*) represents the concentration of the reference signal provided externally to influence the cell behavior. In the above equations, *γ*^e^ and *γ*^j^ are the degradation rate in the environment and in the generic cell of type j (assumed to be the same for all species for the sake of simplicity).

## III. Consortium engineering

In this section, we show that, for the control loop to be effectively closed across the three populations, the parameters characterizing each of the cell populations and the intercellular communication channels must fulfill a set of necessary conditions.

In particular, a set of constraints on the parameters can be derived by analyzing the model equations at steady state, assuming that spatial effects are negligible and the number of cells in the three populations are equally balanced, which implies *η*^j^ = *η*, for all j ∈ {a, d, t}. These assumptions will then be relaxed in the next section where in-silico experiments are carried out also in the presence of cell-to-cell variability and spatio-temporal effects.

### a) Feedback loop pathways

We start by making the realistic assumption that *η* ≫ *γ*^j^, j ∈ {a, d, t}, that is, the signaling molecules diffuse through the cellular membrane faster then they are degraded. Hence, when the reference signal fed to the environment is constant and large enough (i.e. 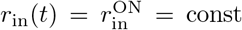) and the Target cells are not active (i.e. *Q_y_* is not expressed), it is easy to verify that at steady state the concentrations of the signaling molecules *R* and *Q_u_i__* reach the same value in every cells, that is, for all j ∈ {a, d, t} we have

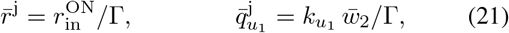

where Γ: = *γ*^e^+*γ*^1^+*γ*^a^+*γ*^d^, and 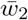 denotes the steady-state value of *w*_2_ when it is fully expressed.

Analogously, for *Q_y_* and *Q*_*u*_2__, when the reference signal *r*_in_ is absent (i.e. 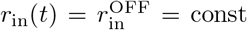) and the Target cells are initially active (i.e. *Q_y_* is expressed), at steady state for all j ∈ {a, d, t} we have

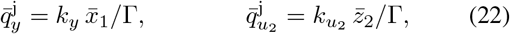

where, similarly, 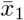 and 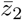 denotes the steady-state values of *x*_1_ and *Z*_2_ when they are fully expressed.

### b) Toggler cells

To guarantee that the Activators and Deactivators implement at steady state the logic functions (1)-(2), it is necessary that only the Activators produce their control signal when the concentrations in the cells of the reference molecule *R* and *Q_y_* are sufficiently high and low, respectively. Therefore, it must hold that

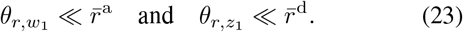

Similarly, when the concentrations of *R* and *Q_y_* are sufficiently low and high, respectively, then, in order that only the Deactivators generate their control signal, it must hold that

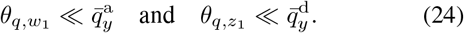

### c) Target cells

In order for the signaling molecules coming from the controllers to toggle the switch within the Targets, the input functions (5) must be such that

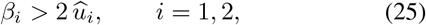

and the concentrations of the quorum sensing molecules within the Targets must be sufficiently high

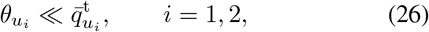

so that the control input is strong enough to trigger the transition from one state to the other and render the toggleswitch monostable.

### d) Parameters’ constraints

Substituting equations (21)-(22) in conditions (23), (24) and (26), we obtain that, at steady state, the Togglers can activate or deactivate the Targets in response to the presence or absence of the external reference signal *r*_in_ (*t*) if the system parameters satisfy (25) and the following conditions are satisfied:

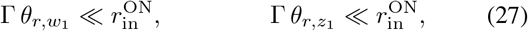

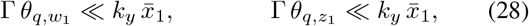

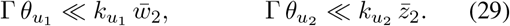

Using similar arguments, it is also possible to obtain lower bounds for the θs, yielding the parameter constraints

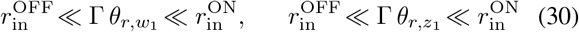

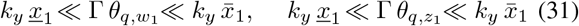

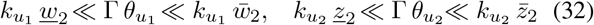

where 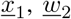 and 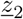 denote the steady-state values of the corresponding species when they are completely repressed.

The previous conditions represent a set of necessary conditions for the consortium to exhibit its desired multicellular control functions.

### Remark

Conditions (30)-(32) depend on steady-state values of *x*_1_, *w*_2_ and *Z*_2_, which in general would need to be estimated in-silico or quantified experimentally. However, at the price of having more relaxed bounds, conditions not depending on these values can be obtained by approximating Hill functions with step functions (i.e. by letting their coefficient *n* → ∞) yielding 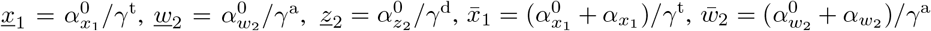 and 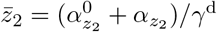.

## IV. IN-SILICO EXPERIMENTS

### A. Agent-based simulations

To validate the effectiveness of our multicellular control design, we implemented a set of in-silico experiments via BSim, a realistic agent-based simulator of bacterial populations [18], [19]. In so doing, we modeled a microfluidics chamber of dimensions 13.3*μ*m × 16.6*μ*m × 1*μ*m and used BSim to take into account cell growth and division, spatial effects, diffusion of the signaling molecules, cell-to-cell variability and geometric constraints. The nominal values of the parameters used in simulations are reported in Table I. They have been chosen similarly as those reported in [7], [15], and satisfying conditions (25) and (30)-(32). Cell-to-cell variability was modeled by assigning a different set of parameters to daughter cells when they split from their mothers. Specifically, each of their parameters, say *μ*, was drawn independently from a normal distribution centered at its nominal value 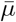 and with coefficient of variation *c*_v_ = 10%.

**TABLE I.** Nominal simulation parameters of the multicellular system

| | [$\mu\text{M}/\text{min}$] | | [$\mu\text{M}$] |
| --- | --- | --- | --- |
| $\alpha_{x_1}^0, \alpha_{x_2}^0$ | 0.005 | $\theta_{x_1}, \theta_{x_2}$ | 2 |
| $\alpha_{x_1}, \alpha_{x_2}$ | 0.1083 | $\theta_{u_1}, \theta_{u_2}$ | 0.5 |
| $\alpha_{z_1,q}^0, \alpha_{z_1,r}^0$ | 0.0348 | $\theta_{r,z_1}, \theta_{q,z_1}$ | 0.5 |
| $\alpha_{z_1,q}, \alpha_{z_1,r}$ | 0.1305 | $\theta_{r,w_1}, \theta_{q,w_1}$ | 0.5 |
| $\alpha_{w_1,q}^0, \alpha_{w_1,r}^0$ | 0.0348 | $\theta_{z_1}, \theta_{w_1}$ | 0.5 |
| $\alpha_{w_1,q}, \alpha_{w_1,r}$ | 0.1305 | | [1/min] |
| $\alpha_{z_2}^0, \alpha_{w_2}^0$ | 0.0016 | $\gamma^t$ | 0.0092 |
| $\alpha_{z_2}, \alpha_{w_2}$ | 0.026 | $\gamma^d, \gamma^a, \gamma^e$ | 0.0230 |
| $\beta_1$ | 0.09 | $\eta$ | 2 |
| $\beta_2$ | 0.016 | $k_y$ | 0.03 |
| $n$ | 2 | $k_{u_1}$ | 0.06 |
| | | $k_{u_2}$ | 0.06 |

Figs. 3 and 4 show the results of a typical in-silico experiment where the Toggler cells (depicted in red and green) successfully flip the Target cells from their active state (depicted in blue) to their inactive state (depicted in black) and vice versa, following changes in the reference signal *r*_in_. The amplitude and the duration of the reference pulse *r*_in_ have been heuristically set to 43 *μ*M and 1140 min, respectively (A more accurate tuning of the pulse can be done by means of other methods, such as in [20].).

**Fig. 3.**
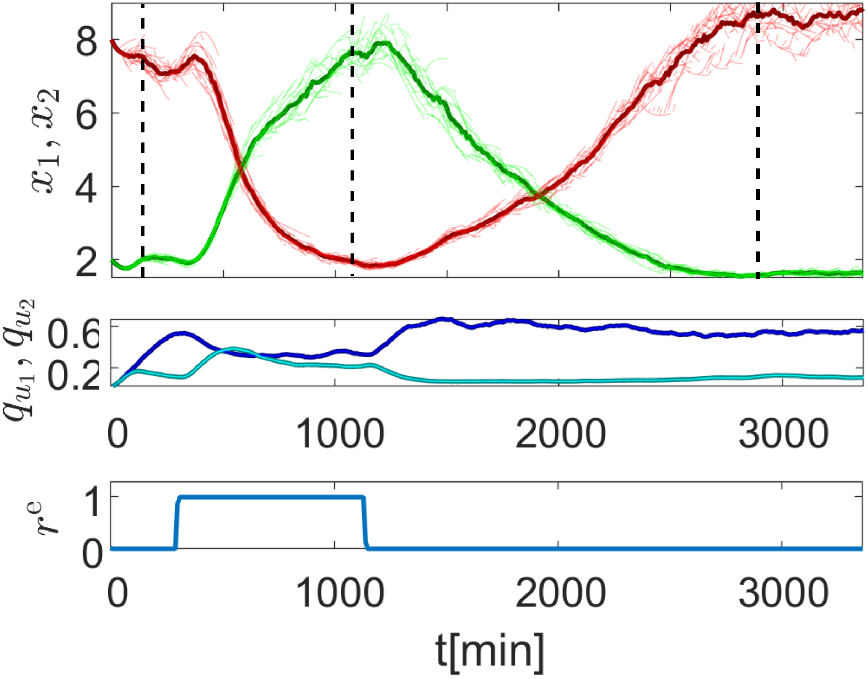
Evolution of the average (thick lines) and single cell (thin lines) values of the concentrations of repressor proteins *x*_1_ (green) and *x*_2_ (red) in the Target population (top panel) when the reference signal *r*_in_ (*t*) is switched from low to high and vice versa. The middle panel shows the average value of the concentrations of quorum sensing molecules 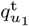 (light blue) and 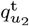 (dark blue) inside the Target cells. The bottom panel shows the value of *r^e^* at the center of the chamber.

**Fig. 4.**
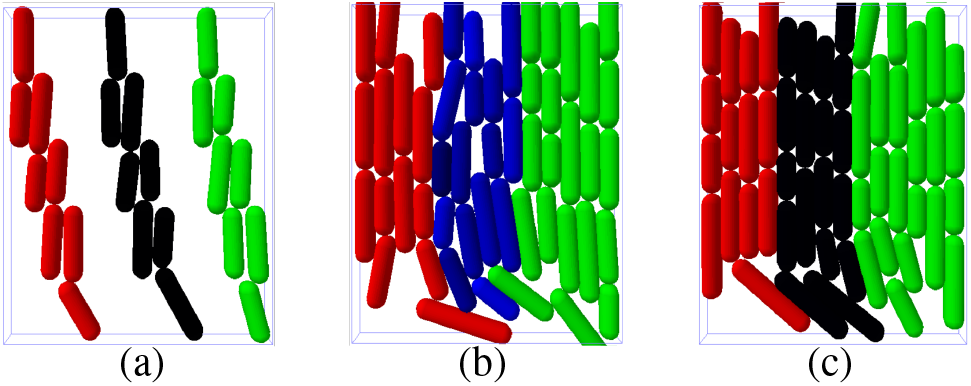
Snapshots of an agent-based simulation at different time instants (highlighted in Fig. 3 with dashed vertical lines). Specifically, panel (a) corresponds to *t* = 120min, panel (b) to *t* = 1100min and panel (c) to *t* = 2800min. Activator cells are shown in green, Deactivator cells in red and Target cells are depicted in blue when they are active and in black when they are inactive.

### B. Robustness to parameter variations

Next, we performed numerical analysis in Matlab, in the illustrative case of no population growth, to evaluate (i) how imbalances between populations affect the operation of the consortium due to poor intercellular communication, and (ii) its robustness to perturbations in the parameters.

Fig. 5 shows the values at steady state of the ratio *x*_1_/*x*_2_ when the Targets are switched OFF (bottom-left panel) and ON (top-right panel), respectively, following the application of the corresponding reference signal *r*_in_, as the ratios of the cell populations in the consortium are being varied. In this scenario, we see that for a wide range of population densities (black region for Deactivators in bottom-left panel, non-black region for Activators in top-right panel of Fig. 5), the Togglers are effectively able to flip the state of Targets.

**Fig. 5.**
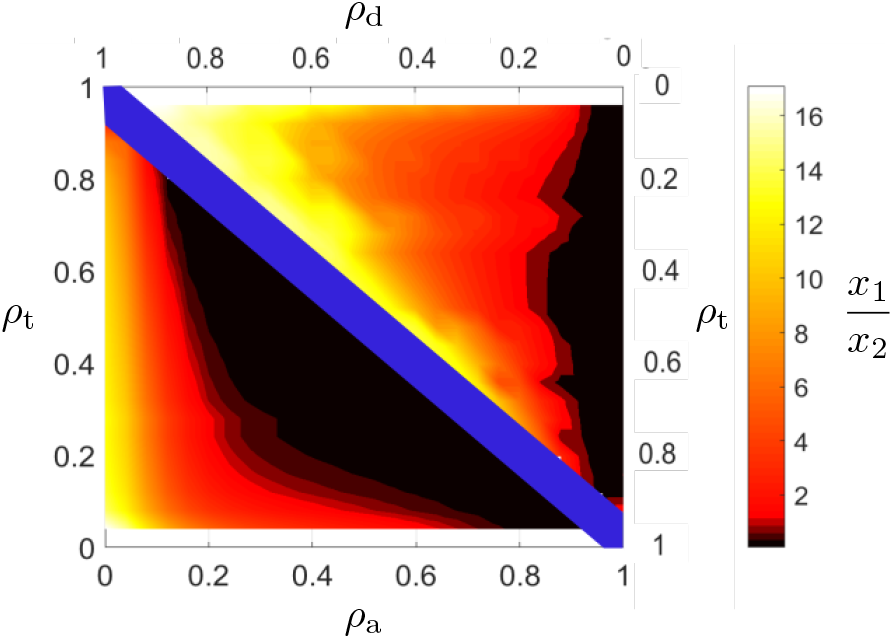
Steady-state value of *x*_1_/*x*_2_ in response to switch commands, 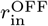 (bottom left) and 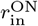 (top right), as the relative ratios of the three populations are changed. Switch commands are applied with the Targets starting from the opposite state. The total cell population in the consortium is set to *N* = 50. For each cell type *ρ*_j_ = *N*_j_/*N*, j = {t, a, d}, is its relative ratio within the consortium, and such that *ρ*_a_ + *ρ*_d_ + *ρ*_t_ = 1.

Finally, we tested robustness of our design when all parameters of Targets, Activators and Deactivators are perturbed from their nominal values. As shown in Fig. 6, even in the presence of a consistent parameter mismatch (*c*_v_ = 0.2), the Togglers are able to activate or deactivate a large fraction of the Targets’ population with the Activators showing better performance of the Deactivators in the parameters’ region we selected.

**Fig. 6.**
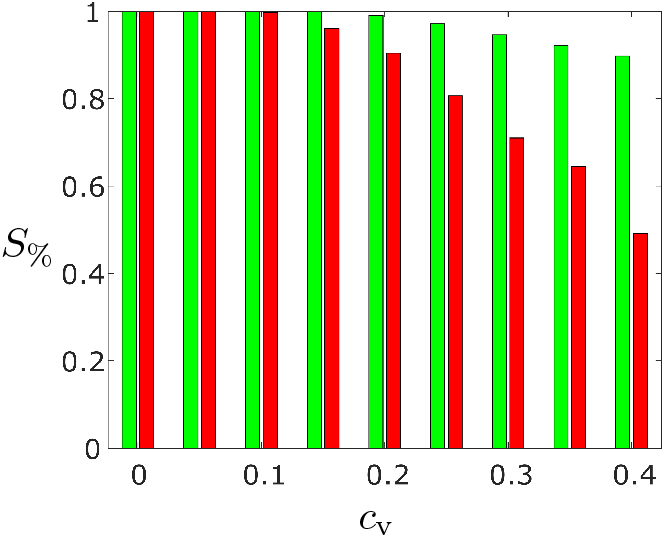
Percentage of successfully switched Targets (*S*_%_) in a balanced consortium (*N*_t_ = *N*_a_ = *N*_d_ = 17) as the coefficient of variation *c*_v_ is varied. The bar plot in red (green) represents the percentage at steady state of Targets that, starting from ON (OFF) state, are turned OFF (ON) following the reference input *r*_in_ being switched to 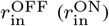. For each value of *c*_v_, the results of 100 simulations were averaged, each obtained by drawing independently all cells’ parameters from normal distributions centered on their nominal values, 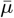, and with standard deviation 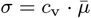.

## V. DISCUSSION AND CONCLUSIONS

We have presented a multicellular control solution to the problem of toggling a memory mechanism in a target cell population. The approach we presented consists in engineering a synthetic microbial consortium of three populations, where two of them act as controllers able to induce activation or deactivation of an inducible genetic switch in the third population, in response to some external reference input. After modeling all the essential components of our design, we discussed some feasibility issues before presenting a careful in-silico investigation of the proposed approach and its robustness to changes in the ratios between the cell populations in the consortium and to perturbation in the parameter values. Our results confirm that the solution we propose is theoretically viable.

The in-vivo implementation of the consortium we propose is currently beyond what is technologically possible but is not unrealistic given that each of the controller population is similar to the comparator implemented experimentally in [21] and the antithetic feedback controller recently presented in [22]. Also, orthogonal communication channels able to set in place the required interaction between the three populations are available and have been tested in [23]. Due to different growth rates of the strains involved, a crucial open problem when constructing synthetic cell consortia is to guarantee stable co-existence and maintain a desired ratio [24], [25] between them to avoid exiting the regions shown in Fig. 5 where the required functions are guaranteed.

We wish to emphasize that a simpler implementation of our design can be obtained by considering a consortium comprising the Target cells and only one population of Activators (or Deactivators), so that the multicellular control strategy is still able to activate initially inactive Targets (or deactivate initially active ones, respectively). With this respect our modular multicellular design is resilient as some functions of the consortium will still be present if one controller population is lost (e.g. becomes extinct or washed out), allowing also for more flexible deployment of the strains in the environment.

A possible future application of the design we propose could be its use for the controlled delivery of active molecules or drugs synthesized by the Targets when they are active. Indeed, the consortium is designed so that the drug or active molecule of interest is only produced and secreted when a specific *reference* chemical signal is perceived by the controller cells with production being stopped when such a reference is removed and the Targets are deactivated. By using a cancer biomarker as the reference signal, the Togglers could then be used to activate the Targets to deliver chemotherapy drugs in situ only when the biomarker is detected in the tissue, providing a multicellular feedback control alternative to the open-loop design proposed in [8]. Also, if the reference signal was linked to the presence of some pollutant in the environment, the controlled activation of the Targets cells could be used to synthesize active molecules for bioremediation when and where they are needed.

## ACKNOWLEDGMENTS

MdB, DF and DS wish to acknowledge support from the European Union’s Horizon 2020 research and innovation programme under grant agreement No 766840 (COSY-BIO). MdB and JO wish to acknowledge support from the University of Naples Federico II Internationalization fund that supported visits and student exchanges within the framework agreement between UPC and the University of Naples Federico II. JO wishes to acknowledge partial support from the Government of Spain through the Agencia Estatal de Investigatiòn Project DPI2017-85404-P and by the Generalitat de Catalunya through the Project 2017 SGR 872.

## References

[1] D. Del Vecchio, Y. Qian, R. M. Murray, and E. D. Sontag, “Future systems and control research in synthetic biology,” Annual Reviews in Control, vol. 45, pp. 5–17, 2018.

[2] J. R. Van Der Meer and S. Belkin, “Where microbiology meets microengineering: design and applications of reporter bacteria,” Nature Reviews Microbiology, vol. 8, pp. 511–522, 2010.

[3] U. Wegmann, A. L. Carvalho, M. Stocks, and S. R. Carding, “Use of genetically modified bacteria for drug delivery in humans: Revisiting the safety aspect,” Scientific Reports, vol. 7, no. 2294, 2017.

[4] D. B. Pedrolli, N. V. Ribeiro, P. N. Squizato, et al., “Engineering microbial living therapeutics: The synthetic biology toolbox,” Trends in Biotechnology, vol. 37, no. 1, pp. 100–115, 2019.

[5] E.-M. Nikolados, A. Y. Weiße, F. Ceroni, and D. A. Oyarzún, “Growth defects and loss-of-function in synthetic gene circuits,” ACS Synthetic Biology, vol. 8, no. 6, pp. 1231–1240, 2019.

[6] P. Bittihn, M. O. Din, L. S. Tsimring, and J. Hasty, “Rational engineering of synthetic microbial systems: from single cells to consortia,” Current Opinion in Microbiology, vol. 45, pp. 92–99, 2018.

[7] G. Fiore, A. Matyjaszkiewicz, F. Annunziata, et al., “In-silico analysis and implementation of a multicellular feedback control strategy in a synthetic bacterial consortium,” ACS Synthetic Biology, vol. 6, no. 3, pp. 507–517, 2017.

[8] M. O. Din, T. Danino, A. Prindle, et al., “Synchronized cycles of bacterial lysis for in vivo delivery,” Nature, vol. 536, pp. 81–85, 2016.

[9] P. Hillenbrand, G. Fritz, and U. Gerland, “Biological signal processing with a genetic toggle switch,” PLoS One, vol. 8, no. 7, 2013.

[10] T. S. Gardner, C. R. Cantor, and J. J. Collins, “Construction of a genetic toggle switch in Escherichia coli,” Nature, vol. 403, p. 339–342, 2000.

[11] J.-B. Lugagne, S. S. Carrillo, M. Kirch, et al., “Balancing a genetic toggle switch by real-time feedback control and periodic forcing,” Nature Communications, vol. 8, no. 1671, 2017.

[12] D. Fiore, A. Guarino, and M. di Bernardo, “Analysis and control of genetic toggle switches subject to periodic multi-input stimulation,” IEEE Control Systems Letters, vol. 3, no. 2, pp. 278–283, 2019.

[13] A. Guarino, D. Fiore, and M. di Bernardo, “In-silico feedback control of a MIMO synthetic toggle switch via pulse-width modulation,” in Proc. of the 18th European Control Conference, 2019, pp. 680–685.

[14] A. Guarino, D. Fiore, D. Salzano, and M. di Bernardo, “Balancing cell populations endowed with a synthetic toggle switch via adaptive pulsatile feedback control,” ACS Synthetic Biology, vol. 9, no. 4, pp. 793–803, 2020.

[15] C. C. Samaniego, N. A. Delateur, G. Giordano, and E. Franco, “Biomolecular stabilisation near the unstable equilibrium of a biological system,” in Proc. of the 58th IEEE Conference on Decision and Control, 2019, pp. 958–964.

[16] C. C. Samaniego and E. Franco, “A robust molecular network motif for period-doubling devices,” ACS Synthetic Biology, vol. 7, no. 1, pp. 75–85, 2018.

[17] J. L. Cherry and F. R. Adler, “How to make a biological switch,” Journal of Theoretical Biology, vol. 203, no. 2, pp. 117–133, 2000.

[18] T. E. Gorochowski, A. Matyjaszkiewicz, T. Todd, et al., “BSim: An agent-based tool for modeling bacterial populations in systems and synthetic biology,” PLoS One, vol. 7, no. 8, p. e42790, 2012.

[19] A. Matyjaszkiewicz, G. Fiore, F. Annunziata, et al., “BSim 2.0: an advanced agent-based cell simulator,” ACS Synthetic Biology, vol. 6, no. 10, pp. 1969–1972, 2017.

[20] A. Sootla, D. Oyarzún, D. Angeli, and G.-B. Stan, “Shaping pulses to control bistable systems: Analysis, computation and counterexamples,” Automatica, vol. 63, pp. 254–264, 2016.

[21] F. Annunziata, A. Matyjaszkiewicz, G. Fiore, et al., “An orthogonal multi-input integration system to control gene expression in Escherichia coli,” ACS Synthetic Biology, vol. 6, no. 10, pp. 1816–1824, 2017.

[22] S. K. Aoki, G. Lillacci, A. Gupta, et al., “A universal biomolecular integral feedback controller for robust perfect adaptation,” Nature, vol. 570, pp. 533–537, 2019.

[23] N. Kylilis, Z. A. Tuza, G.-B. Stan, and K. M. Polizzi, “Tools for engineering coordinated system behaviour in synthetic microbial consortia,” Nature Communications, vol. 9, no. 2667, 2018.

[24] D. Salzano, D. Fiore, and M. di Bernardo, “Ratiometric control for differentiation of cell populations endowed with synthetic toggle switches,” in Proc. of the 58th IEEE Conference on Decision and Control, 2019, pp. 927–932.

[25] X. Ren, A.-A. Baetica, A. Swaminathan, and R. M. Murray, “Population regulation in microbial consortia using dual feedback control,” in Proc. of the 56th IEEE Conference on Decision and Control, 2017, pp. 5341–5347.

